# Gender balance in time-keeping at life science conferences

**DOI:** 10.1101/267120

**Authors:** Petra Edlund, Erin M. Tranfield, Vera van Noort, Karen Siu Ting, Sofia Tapani, Johanna L. Hoog

## Abstract

Scientific conferences are biased against women when selecting speakers, but recent studies showed a positive trend where the ratio of invited female speakers had increased. We have recorded speaking times at 11 different conferences during 2016-2017 to analyze which factors influenced a speaker’s time keeping. Men and women went over time 47% and 41% of the talks, respectively. In a regression analysis, it was found that the most important factors that influences how long a presenter spoke was 1) their allocated time, 2) their career stage and 3) the level of time keeping enforcement. It was also found that gender and the size of the conference contributed significantly towards the speaker’s timing. Male speakers exceeded their allocated time more frequently than female speakers, especially at large conferences (60% vs 49%). Since conferences are an important arena for science dissemination this might have a negative impact on female scientist’s careers.

## Introduction

The last few years have seen an increased attention to the lack of women representation in faculty positions in the natural science departments of universities and institutes around the world (1–3). What makes this issue even more surprising is that a majority of students graduating with bachelors or masters in science, technology, engineering and mathematics (STEM) are women (4). Almost half of all the European PhD students are female (5) and in life sciences in particular, 55% of the PhD students were female in the USA 2015, an equivalent number for Europe could not be found. At postdoctoral level the number of female researchers’ decreases and it drops dramatically in leadership positions (1). Therefore, something happens during their academic career that deters women from either pursuing a continued career in scientific research, or from achieving success in their research, and are therefore unable to progress. This is commonly denoted as “the glass ceiling”(1). This glass ceiling is a waste of useful resources and talent both for the research community and society itself.

A scientific career in academia demands varied tasks such as teaching, designing and carrying out research, reading literature, preparing grant applications and engage in scientific outreach activities. But the most determinant in their career progression is to communicate their research findings. To achieve this, scientists write and publish academic papers and present at scientific conferences. An academic’s scientific skill is largely measured by their metrics (i.e. number of articles, their impact factor and citations) of the articles they produce (6). The other way is to highlight and promote the research that has been done and to present their findings at a scientific conference and university colloquia. The participation in these scientific events create visibility for researchers and open networking opportunities, with potential to kick start projects or even obtain jobs. A series of studies has shown that in these situations, there is a bias against selecting women to be scientific speakers (7–12).

However, the trends are positive with American Society for Microbiology General Meeting declared speaker gender balance in 2015 (13), and around 40% female invited speakers at major virology conferences in 2017 (7). However, we do not know how gender balanced the conferences are in terms of the actual time being spoken. In this study, we compare the gender balance of speakers to the genders of the conference attendees, and measure how much actual speaking time male and female speakers occupied with respect to their allocated times. Furthermore, we compared the compliance to the allocated time between PhD students, postdocs and PIs to investigate this problem.

## Results

### Number of female speakers correlates well with the number of female participants at the studied conferences

Previous studies have shown a bias against women when selecting speakers for scientific conferences. We quantified the gender of speakers at 11 conferences; both the number of talks given by women and men respectively, but also their allocated and actual talk time in minutes (Figure 1).

**Figure 1:**
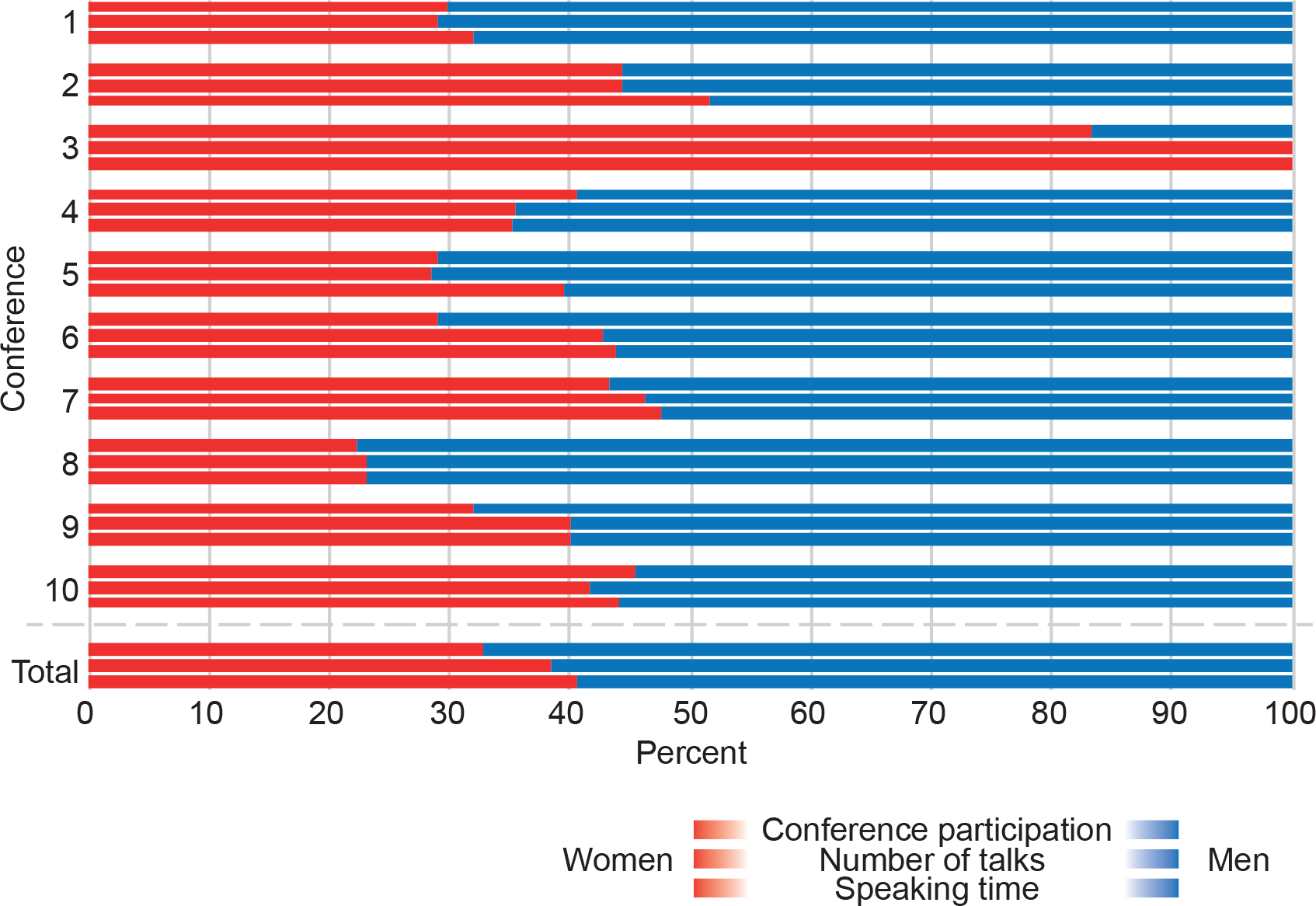
Women were well represented as speakers when compared to their rate of attendance. The graph shows the gender distribution in ten of studied conferences, where women’s contributions are visualized in red and men’s in blue. For each conference, we show gender distribution in attendance, number of talks and the actual time that they spoke.

To compare the number of talks given/talk time of each gender with the gender distribution of conference participants, the conference organizers were asked to provide the numbers of participants, and their respective gender (10 organizers provided this). In total only 33% of the 3487 conference participants were female. However, women gave 38% of the scientific presentations. Since there is a larger population female young scientists (4) one could imagine that the female speakers were early career scientists who generally get shorter talks. This was not the case though, with the average talk length not differing between men (22 min; n=189) and women (23 min; n=118), and women speaking 40% of the total presentation time. There might even be a slight preference for female speakers since they were selected to speak slightly more than they were attending at 8 out of 10 conferences.

Conference 3 is an outlier, where most of the participants and all the speakers were female. This should be highlighted as an example for future discussion since this was the only regional conference of the 11 conferences that was studied.

We conclude that the studied conferences have a low female participation in relation to representation within the field, but that this participation is fairly represented in the conference speaker program. However, if one gender is better or worse at sticking to time, this organizational gender balance could still be disrupted by a real gender bias. Therefore, the actual time talking was measured and studied in relation with the scheduled time.

### Actual talk time showed differences between genders

Speakers were timed from when they got the word until end of their final slide. Discussion time was not measured. In absolute terms, male speakers did go over time more often than women (Figure2A; 47% and 41% respectively; n=189 male and 118 female talks). A linear regression statistical analysis (Table 1) confirmed that the difference between male and female talking has a significant difference when all other measured factors are accounted for. Generally, male speakers went over time more often than females at the measured conferences. The distribution of timekeeping between men and women are demonstrated in figure 2B-C and show close to a normal distribution. The mean value for used time (percentage of allocated) for men and women speaking time was similar but slightly higher for men (99.5 for men and 97.5 for women).

**Figure 2:**
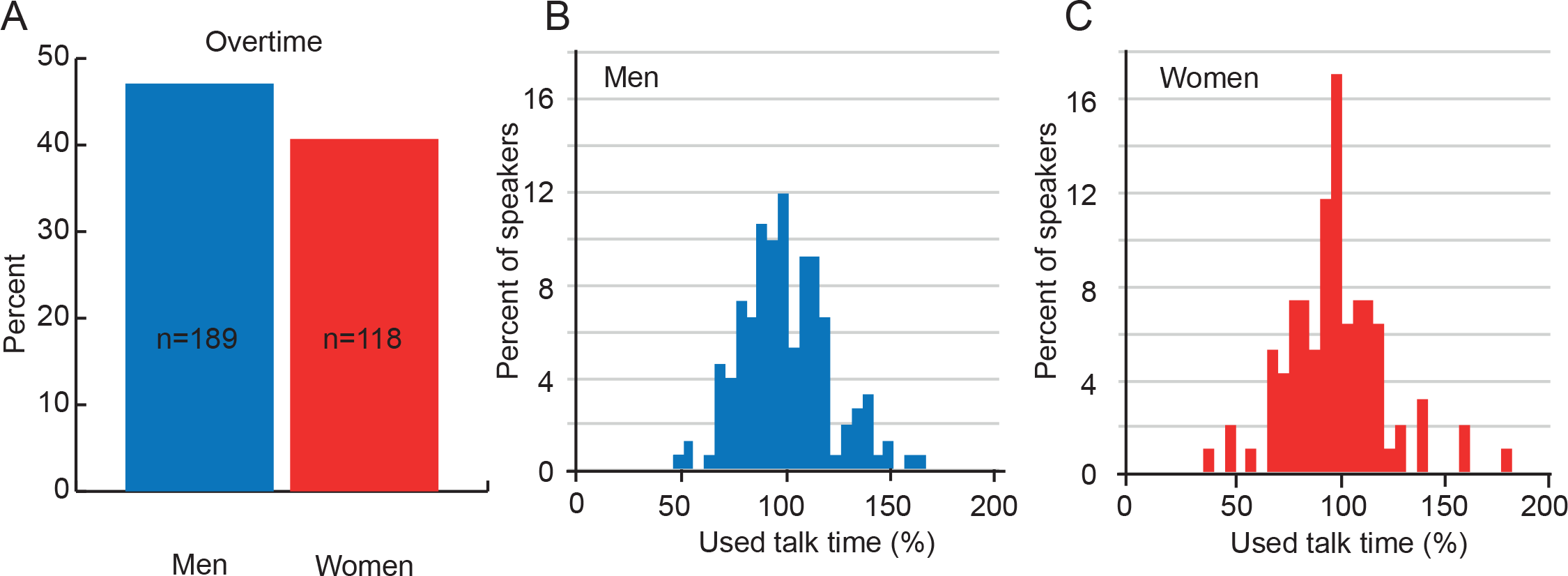
Both men go over time slightly more often. The graphs show A) The percentage of the speakers exceeding their allocated time divided by their gender (blue for men and red for women). B) Show the histogram of male speakers used time as percentage of their allocated time. C) Show the equivalent to B but for female speakers.

**Table 1:**
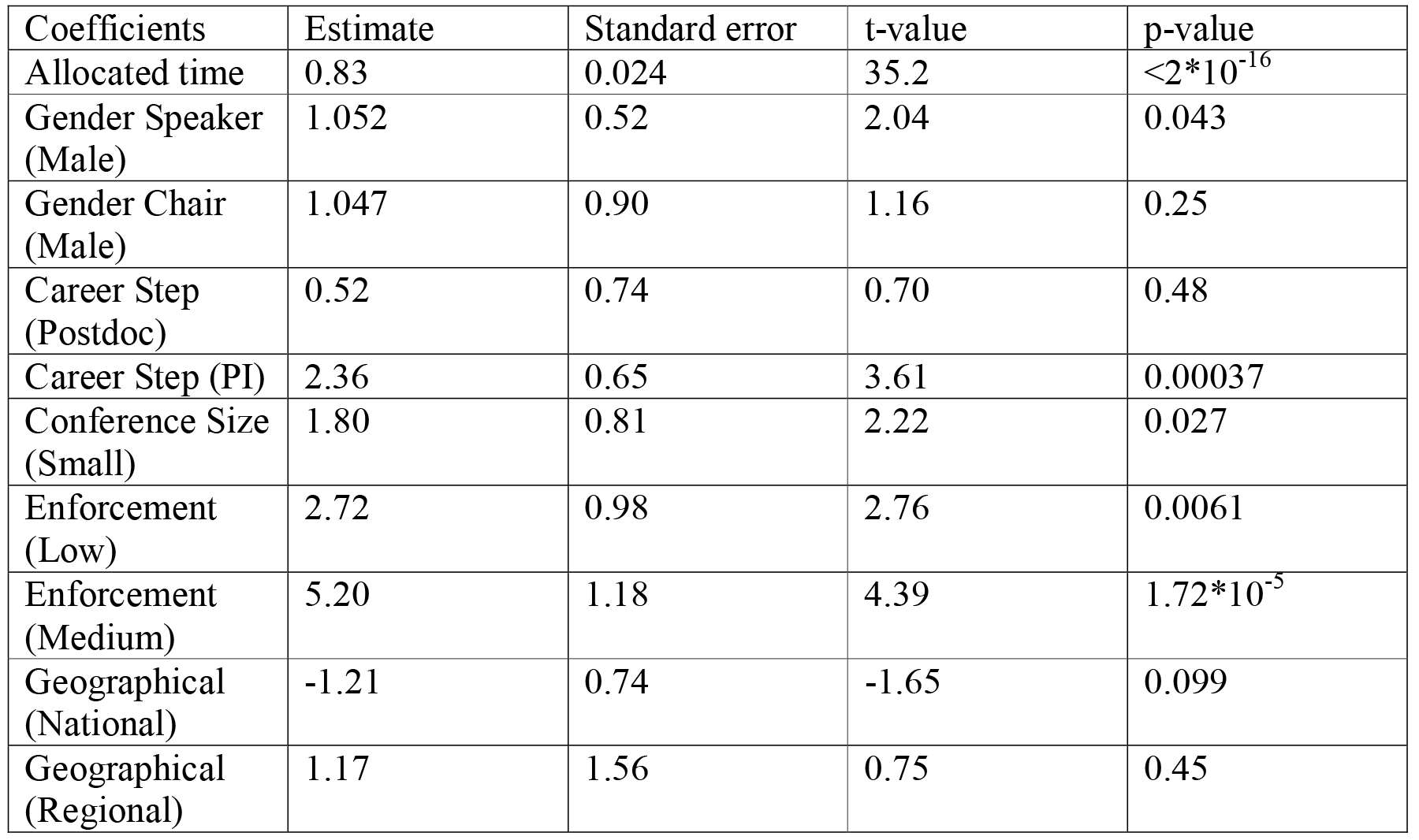
Regression analysis of measured time, model coefficients and their corresponding statistics with tests for coefficient being different from 0.

### PhD students are better at keeping to allocated times than postdocs and PIs

The career step of speakers was also noted to determine the time keeping differences between PhD students early in their career, with later career steps such as postdoctoral or Principal Investigator(PI) speakers. All speakers were divided into 54.7% being PIs, 17.8% postdocs and 27.5% PhDs (Figure 3A; n=236). Only 35.7% of the postdoctoral and PI speakers were female whereas the distribution among PhD students were more equal between genders with 46.2% females (Figure 3B).

**Figure 3:**
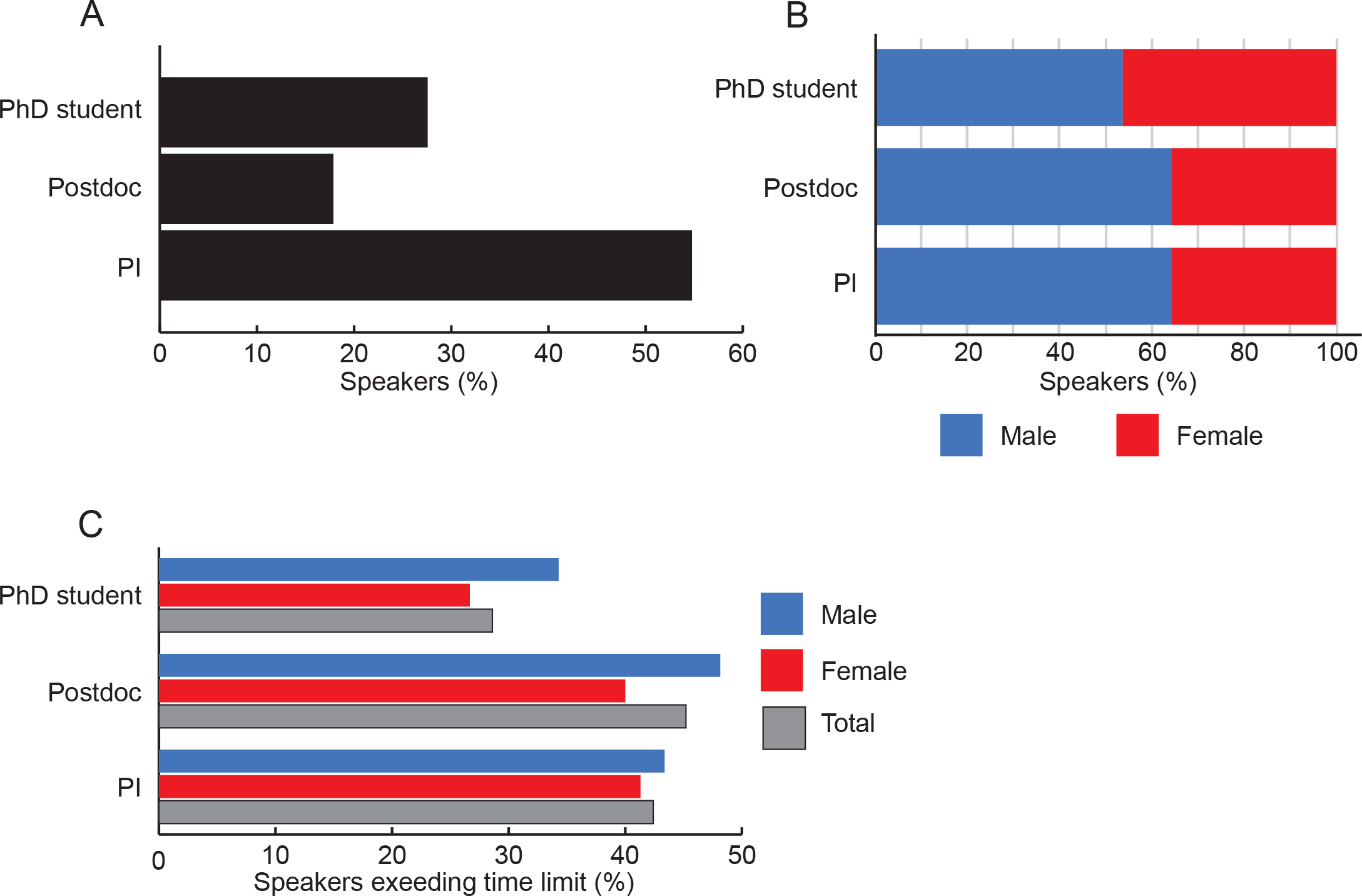
PhD students are best to adhere to time. A) The career stage of the speakers of both genders shows that in these conferences the PIs were the most common speakers. B) The gender distribution of the speakers is only among the PhD student speakers. Postdocs and PIs have a majority of male speakers. C) The percentage of speakers, from the three career steps, that exceeded their allocated time divided by gender and in total. Female PhD students are best at time keeping, and male postdocs are worst. Male in blue, female in red and together in grey.

In general, PhD students were best at time keeping with only 28.6% going over time compared to postdocs 45% and PIs 43% (Figure 3C). The group that was absolutely best at keeping to time was female PhD students (74% on time). In general, postdocs went over time more frequently, with females being 40% overtime and males as high as 48% of the talks. The difference between female and male PIs exceeding time was smaller with 41 % of the females exceeding time and 43% of the males. However, the statistical analysis showed that the PIs exceeded allocated time with a high significance when all other factors were accounted for.

### Male speakers go over time in 60 % of their talks at large conferences

One further aspect of the study was to include the genders of the session chair and evaluate the effect on speakers exceeding the allocated time. Male and mixed chairs had more speakers going over time than female chair had (Figure 4A). Male speakers exceed the time to a greater extent when the chair was male and the females spoke longer when there were two session chairs (one of each sex). Since big part of the chairs were of mixed chair it is hard to draw any conclusion depending on the gender of the chair, and the statistical analysis showed no significant influence on speaking time by the gender of the chair.

**Figure 4:**
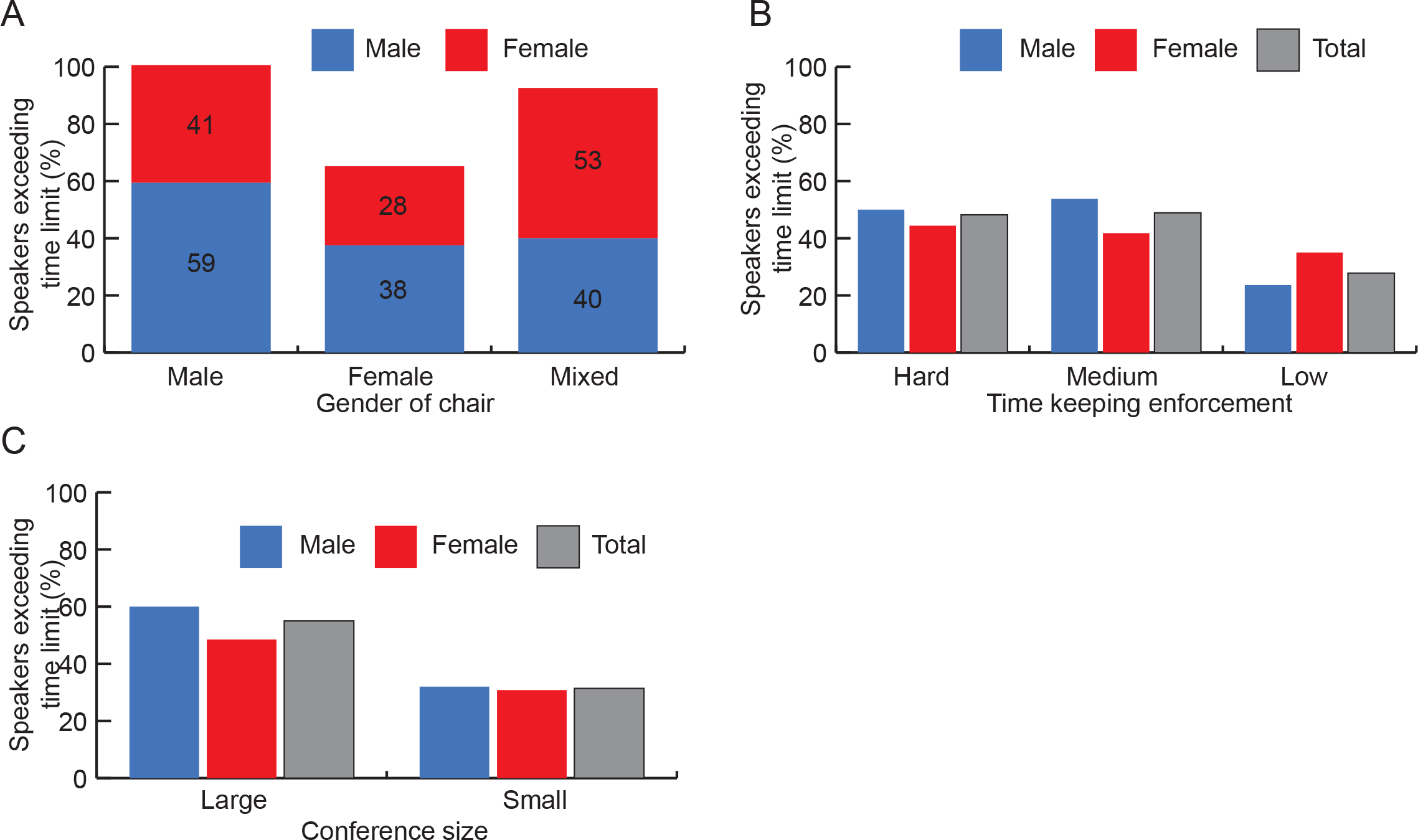
The chair gender, strictness of time keeping enforcement and conference size influenced time keeping. A) The percentage of males (blue) and females (red) speakers exceeding their allocated time related to the gender of the chair at their session. B) The percentage of speakers exceeding their allocated time in relation to the level of time keeping enforcement. Total is shown in gray, males in blue and females in red. C) Show the percentage of speakers exceeding their allocated time in large (>150 participants) versus small (<150 participants) conferences. Total is shown in gray, males in blue and females in red.

The time-enforcement activities showed no major success in keeping speakers on time. Surprisingly the conferences with no or very low enforcement to keep on time had the fewest speakers exceeding time (around 30%) and conferences with hard and medium enforcements speakers went over time close to 50% of the talks (Figure 4B). In addition, the statistical analysis showed that having a medium or low enforcement on keeping to time had a significant impact on speakers going over time. When comparing “large” conferences (>150 participants) to “small“(<150 participants) it was clear that speakers were more likely to exceed their allocated time at large conferences with 55% of all speakers exceeding their time compared to 31% at small conferences (Figure 4C; n =170 speakers at large conferences and n = 137 speakers at small conferences). Although the major part going over time at large conferences was the contribution of male speakers with as much as 60% going over time whilst a smaller share (48%) of the female speakers went over time. At small conferences, around 30% went over time independent of gender of speakers.

## Method

### Study set-up

Social media was used to recruit co-authors to note down speaker times at life science conferences. The Facebook status showed preliminary data collected at one conference and attracted comments from 17 friends who either contributed with comments on how such a study could be performed (6 males and 1 female) and 11 friends (3 men and 8 women) who wanted to contribute measurements. The 11 people who wanted to collect data was sent a template excel file to fill in during the conference (supplemental data). 5 women returned with measurements and one additional woman contributed (recruited by talking about the project). A Twitter was used to try to expand the measurements but yielded no interest.

Each co-author was also asked to note how time keeping was enforced, the gender of the chair(s) and the kind of conference they were attending (regional, national or international). Altogether, we collected time-keeping measurements from 11 conferences between May 2016 and October 2017. The geographical distribution was the following: 2 conferences in Sweden, 2 in Norway, 2 in Germany and 1 each in UK, France, Spain, Holland and the USA. 7 conferences were international, 3 national and one regional. Each speaker was timed until the end of their final slide. The duration allocated for each talk was taken either from the program or from the organizers. The discussion time was not measured. When applicable the career position of the speaker was noted. The measuring routine was kept discrete to avoid biased measurements. The measured time was divided by the allocated time to get the used time in percent for each measurement. The measured data was analyzed using a linear regression (ANOVA) that points out the variables that has a significant effect on the measured talking time (Table 1). A model was fitted to the data with measured spoken time as dependent variable and gender of speaker, gender of chair, allocated time, career step, enforcement, conference size and geographical placement of conference as explanatory variables. A p-value for testing the coefficients in the model being different from 0 less than 0.05 was considered statistically significant.

### Enforcement of time-keeping

The attention paid to making speakers sticking to time varied between conferences.

#### High enforcement

Two conferences were held in a venue that applied a traffic light system. The traffic light started off green, switched to yellow when the end of the speaker time came close and red when the time was out. The red light was further enhanced with the light in the auditorium being switched on and the person responsible for audio visions coming up to the stage. We judge this the most tightly controlled time keeping.

#### Medium enforcement

In four conferences, the chair gave a wave when the speaker had a short time left, and then stood up when their talk time was out. Timekeeping at one conference was done with a big clock in the back of the room, visible by the speaker indicating the remaining time.

#### Low Enforcement

Four conferences had no time-keeping enforcement at all.

## Discussion

Although the difference in timekeeping between genders was not immense. There was a significant difference when all other factors were accounted for, showing that men go over time more often than women. With both genders exceeding time in more than 40% of the talks, it reveals a general disregard for time keeping. The reasons for exceeding the allocated time can be numerous, inadequate preparation, nervousness or tendency to over-value the importance of one’s own research. As we all know, the real results of speakers exceeding time is less time for discussion, later speakers getting their time cut or to distortion of the conference schedule and should therefore, in our opinion, be strictly avoided. We suggest that conference organizers set a time for speaking, followed by a defined time for questions and discussion. If the speaker goes over time, the chair should only allow the discussion to fill the time remaining, and if the speaker talk exceed their total time, that they are told to stop. Another way forward would be to shut off the microphone after the speaker is exceeding their time with 10%, no exceptions. This would encourage speakers to prepare their lectures and time them in advance.

The general trend is positive for women in the aspect that they are given slightly more talks and speaking time than their representation at the conferences. This follows well with the past years positive trend of increasing number of selected female speakers, and is probably a positive response to the previous highlighting of the problem. With conference organizers being more aware, active measures are being taken to prevent the continuation of bias against women, for example initiatives such as the website BiasWatchNeuro which displays the number of female speakers at neuroscience conferences and compares that to the base rates of women found in that field (14). To aid the easy identification of possible female speakers, several lists of prominent women in different research fields has been composed (e.g. “Academia Net” (15) and Anne’s List (16)). To include one or more women on the organizing committee has shown to have a strong positive effect on the number of female speakers of that conference (10, 17). A helpful checklist for conference organizers has been published(18). However, the conference speakers are still far from half women, and if one compares to their presence in the field, a fair number of female speakers should probably be higher than 50%. Why is there not a female participation corresponding to the actual portion of women active in the field of life sciences? One might expect a higher fraction of female participants in these life science conferences, since the field has a large proportion of women. Because of the nature of our study, measuring individual speaking times, the number of studied conferences is rather limited (n=11). It would be interesting to make a larger study of gender distribution at life science conferences to investigate if our low female attendee numbers persist. If so, the reasons behind the lower female attendance needs to be examined in detail to be able to design solutions so that women can display their results to the scientific community as often as men do. One factor seems to be travel, since the one regional conference in this study had a very strong female presence. Therefore, maybe one improvement would be to increase funding for more regional conferences or web-based conferences.

The majority of speakers were PIs, which can be expected since they are more experienced scientists, usually presenting a wider scope of scientific results. Successful keynote speakers are also a major strategy to attract participants to conferences by organizers. Both at the PI and postdoc level there is a substantially larger fraction of male speakers (64.7%). However, the gender balance between PhDs was more even with 46% females. This probably mirrors the female gender distribution within the field to some extent, having around 50% female PhDs (5), fewer postdocs and substantially fewer female PIs. This shows that the lack of female at higher career steps is compensated for, at conferences, by letting a bigger fraction of female PhD students speak. Thus, initiatives in trying to find and promote successful female PIs is not working as efficiently as one would hope. Even if this shows that younger women scientists get the opportunity to show their science, it also shows the lack of female role models and research leaders. This can be a harmful factor for the whole scientific environment, as it is a direct visualization of the glass ceiling phenomenon which might deter young female scientists from pursing a long-term career in research. More visible successful women in academia could lead to more women pursuing a career in life science research and research in general. One conference is an extreme case where all the speakers and more than 80% of the attendees were female. This was the only regional conference in our study, and could be indicative of that increasing the number of small, regional conferences would increase the female participation in life science conferences.

It makes a difference in which career stage a speaker is. PhD students, and in particular female PhD students, go over time most seldom. This is surprising since PhD students are the least experienced lecturers. It can be due to better preparation for talks by younger speakers and it shows that talks can indeed be prepared to the extent where it more often fits into the allocated time. It is interesting to note that the strong gender difference in time-keeping ability of PhD students does not persist in the postdoctoral and PI speakers. One potential explanation for this is that there is selection against time conscious (female) PhD students in academia. In agreement with this, male postdocs is the group exceeding their allocated time most frequently and they also continue their career in research to much higher extent than their female counterparts.

Finally, we evaluated if the gender of the chair, the degree of time keeping enforcement used and size of the conference influenced speakers remaining within their allocated time. The effect of gender of the chair was hard to analyze since there were many mixed gender chairs. The time enforcement does not seem to yield the desired effect and having a hard enforcement yielded only more speakers to exceed their time. Comparing the size of the conference also showed that both genders but especially males are more prone to exceed time at large conferences. These conferences are the biggest arenas to promote yourself and your research and that male speakers go over their allocated time here can contribute to limit female visibility in the life sciences field. As a consequence, this might have a negative impact on the scientific careers of women. On the other hand, one might see it as a sign of professionalism when speakers adhere to time restrictions and maybe speakers that exceed their allocated time are actually not reaping any benefits from this behavior.

None of the organizers reported on non-binary participants and therefore it is not included in the study. Although it should be considered by conference organizers that not everyone identifies as male or female.

Together this study suggests that a conference would have the highest chances of staying to the planned schedule if speakers were preferentially (female) PhD students, had female chairs of sessions, less than 150 participants and no direct enforcement of time keeping at all. The challenges in minimizing gender bias at conferences lies in inviting female PIs to share their science to a greater extent, find ways to get ratio female attendees to reflect their presence in the field and develop methods to enforce speakers to stick to their allocated time. The authors of this paper would also like to highlight the comment of one conference participant: “Personally, I am in favor of a trap door - you go over time and it opens”, illustrating the frustration in the field for the disrespect of allocated speaker times.

## Acknowledgements

We thank all conference organizers that shared participants genders and other important information to us for this study. We also thank Pernilla Wittung-Stafshede for reading and advising on the manuscript. This study was performed without any direct funding as a pro bono collaboration between the all-female scientists authors.

